# Looking for Tumor Specific Transcription Factors. Study of Promoters in Silico

**DOI:** 10.1101/2021.11.11.468214

**Authors:** K.N. Kashkin

## Abstract

This study supplements earlier received experimental data using modern databases. Previously tumor-specific activity of several human native and chimeric promoters was demonstrated. Here we compared tumor-specific promoters with promoters of housekeeping genes by the presence of recognition profiles for transcription factors in DNA sequences of the promoters. A number of transcription factor recognition profiles have been identified, the presence of which in promoters may indicate the tumor specificity of the promoters. Transcription factors which may directly regulate promoters of genes involved in cell proliferation and carcinogenesis were revealed by pathway analysis. The results of the study may help in studying the peculiarities of gene transcription in tumors and in the search for or the creation of tumor-specific promoters for cancer gene therapy.

## INTRODUCTION

The functioning of a eukaryotic promoter is a complex process that includes the interaction of the cis- and trans-regulatory elements of the promoter with transcription factors (TF), enzymes of the transcription complex, and regulatory RNAs [1]. Epigenetic mechanisms, such as conformation changes in chromatin, DNA methylation, histone modifications [2], as well as physicochemical processes associated with phase separation [3] play essential roles in the regulation of transcription. However, binding of TFs to the specific sequences in DNA and the interaction between factors are the basis of the whole regulation of promoter activity.

In total, more than 1600 human TFs are currently known, their number is constantly growing. For a significant part of them (more than 500 factors) the sequences of binding sites in DNA and their exact role are unknown [4]. An essential feature of TFs binding sites is their degeneracy and cross affinity of different TFs for the same sequence [5]. This provides probabilistic and competitive regulation of transcription.

In transformed cells, the peculiarities of gene transcription along with the rearrangement of metabolism determine the ability of tumors to grow, avoiding control by the surrounding tissues and the organism. Therefore, the study of these processes is of theoretical and practical importance for oncology. Some known TFs are considered diagnostic markers of tumors or their metastases. Thus, CDX2 is a sensitive marker of colorectal adenocarcinoma, TTF1 is a significant marker for lung adenocarcinoma, PAX8 can serve as a marker of gynecological tumors, etc. [6]. Therefore, the study of TFs, in particular, the presence of their recognition profiles in tumor-specific promoters, remains relevant.

Earlier, we cloned a number of promoters of genes regulating cell proliferation, which exhibit tumor-specific activity [7, 8] (Table 1). In this work, we compared these promoters as well as the chimeric promoters obtained from them [9] with a set of housekeeping gene promoters in terms of the composition of the known TF recognition profiles.

**Table 1.** The promoters included in the study

| Promoter | EPD index |
| --- | --- |
| *Tumor-specific promoters* |  |
| *BIRC5* | FP024000 |
| *CDC6* | FP023205 |
| *CKS1B* | FP001855 |
| *MCM2* | FP005950 |
| *PLK1* | FP021553 |
| *POLD1* | FP026208 |
| *TERT* | FP007749 |
| *PCNA Promoter* |  |
| *PCNA* | FP026732 |
| *Chimeric promoters* |  |
| *CH2-CH26* | Ref. [9] |
| *Promoters of housekeeping genes* |  |
| *ACTB* | FP010679 |
| *ALAS1* | FP005471 |
| *B2M* | FP020416 |
| *GAPDH* | FP017185 |
| *GUSB* | FP011089, FP011090 |
| *HMBS* | FP016933, FP016934 |
| *HPRT1* | FP029346, FP029347 |
| *PGK1* | FP029028 |
| *PPIA1* | FP011011 |
| *RPL13A* | FP026153 |
| *RPLP0* | FP018538, FP018539 |
| *SDHA* | FP007723 |
| *TBP* | FP010603, FP010604 |
| *TFRC* | FP006502 |
| *TUBB* | FP009374 |
| *YWHAZ* | FP012696, FP012697, FP012698 |

We believe that such an analysis will help to understand the mechanisms of tumor-specific gene transcription better and will open up new possibilities for creating artificial promoters and genetically engineered vectors used in gene therapy of tumors.

## MATERIALS AND METHODS

The sequences of human promoters were obtained from the EPD database (Eukaryotic Promoter database, https://epd.epfl.ch/EPDnew_database.php) [10] at coordinates [−499; +100] relative to the transcription start point (Table 1). The CiiiDER program [11] and the JASPAR2020 database of transcription factor profiles (http://jaspar.genereg.net/) [5] were used to search and compare TF profiles in the promoters. The expression of TFs in tumors is listed by the open GEPIA2 database (http://gepia2.cancer-pku.cn/). The prognostic value of TFs in tumors was determined according to The Human Protein Atlas (https://www.proteinatlas.org/). The Pathway Commons resource (http://www.pathwaycommons.org/) [12] was used to build regulatory networks.

## RESULTS

### Profiles prevailing in tumor-specific promoters

We have previously shown that relatively short (hundreds of nucleotides in size) promoters of a number of genes (Table 1) involved in the regulation of proliferation have tumor-specific activity [7, 8]. It is known also that the most important regulatory elements are concentrated in the region up to –500 base pairs from transcription start sites (TSS) in the studied promoters [8, 13, 14]. Therefore, we selected seven DNA sequences which corresponded to previously investigated tumor-specific promoters from the database of eukaryotic EPD promoters with coordinates [−499; +100] with respect to the TSS. For comparison, we selected 23 promoters of housekeeping genes of the same length from the EPD database (Table 1). We compared tumor-specific promoters with housekeeping gene promoters in terms of the composition of TF recognition profiles using the CiiiDER program [11] (enrichment function) and the JASPAR2020 transcription factor profile database (http://jaspar.genereg.net/) [5]. The number of promoters containing the corresponding TF profiles in both groups was compared, and the results were assessed by the Mann-Whitney test. The full results of the comparison of the promoters are presented in Supplementary Table S1. Table 2 and Supplementary Table S2 show the TF profiles that are significantly (p<0.05) prevailing in tumor-specific promoters in comparison with the promoters of housekeeping genes. These profiles are hereinafter referred to as conditionally tumor-specific factors. The most saturated promoters with the TFs profiles of this group are MCM2 (13 profiles), CKS1B (10 profiles), and PLK1 (9 profiles) promoters (Supplementary Table S3). More than half of tumor-specific promoters contain profiles of the factors SREBF2, ZNF75D, Zfx (each in six promoters), RUNX2 and ETS2 (each in five promoters) and Creb3l2 (in four promoters) (Table 2). For three TFs, it was possible to identify preferential zones of location in the promoters (Fig. 1). SREBF2 profiles in 4 out of 7 promoters are located in the region [−458; −384] relative to TSS on the noncoding DNA strand. ZNF75D profiles in four promoters occupy the region [−253; −184] relative to TSS also on the non-coding DNA strand and in three promoters near TSS [−24; +30] on the coding DNA strand. RUNX2 profiles are located on the coding strand in the region [−414; −247] in 4 promoters.

**Table 2.** Recognition profiles of transcription factors which are more frequent in tumor-specific promoters than in promoters of housekeeping genes*

| **Transcription factor** | **JASPAR 2020 nomenclature** | **Class** | **Family** | **Tumor-specific promoters that contain the profile** | **Number of housekeeping promoters containing the profile\*** |
| --- | --- | --- | --- | --- | --- |
| RUNX2** | MA0511.2 | Runt domain factors | Runt-related factors | *POLD1, BIRC5, PLK1, CDC6, CKS1B* | 4 |
| Creb3l2** | MA0608.1 | Basic leucine zipper factors (bZIP) | CREB-related factors | *MCM2, PLK1, CDC6, TERT* | 2 |
| SREBF2** | MA0596.1 | Basic helix-loop-helix factors (bHLH) | bHLH-ZIP factors | *MCM2, POLD1, BIRC5, PLK1, CKS1B, TERT* | 7 |
| ETS2** | MA1484.1 | Tryptophan cluster factors | Ets-related factors | *MCM2, PLK1, CDC6, CKS1B, TERT* | 5 |
| CENPB | MA0637.1 | CENP-B |  | *MCM2, CDC6, CKS1B* | 1 |
| HEY1 | MA0823.1 | Basic helix-loop-helix factors (bHLH) | Hairy-related factors | *MCM2, PLK1, TERT* | 1 |
| HEY2 | MA0649.1 | Basic helix-loop-helix factors (bHLH) | Hairy-related factors | *MCM2, PLK1, TERT* | 1 |
| RARA::RXRG | MA1149.1 | Nuclear receptors with C4 zinc fingers::Nuclear receptors with C4 zinc fingers | Thyroid hormone receptor-related factors (NR1)::RXR-related receptors (NR2) | *MCM2, POLD1, CKS1B* | 1 |
| SREBF1(var.2) | MA0829.2 | Basic helix-loop-helix factors (bHLH) | bHLH-ZIP factors | *MCM2, BIRC5, PLK1* | 1 |
| ZNF75D** | MA1601.1 | C2H2 zinc finger factors | More than 3 adjacent zinc finger factors | *MCM2, POLD1, PLK1, CDC6, CKS1B, TERT* | 8 |
| Zfx** | MA0146.2 | C2H2 zinc finger factors | More than 3 adjacent zinc finger factors | *MCM2, POLD1, BIRC5, CDC6, CKS1B, TERT* | 9 |
| HOXD11 | MA0908.1 | Homeo domain factors | HOX-related factors | *POLD1, CKS1B* | 0 |
| KLF11 | MA1512.1 | C2H2 zinc finger factors | Three-zinc finger Kruppel-related factors | *POLD1, BIRC5* | 0 |
| MEF2A | MA0052.4 | MADS box factors | Regulators of differentiation | *MCM2, BIRC5* | 0 |
| RORB | MA1150.1 | Nuclear receptors with C4 zinc fingers | Thyroid hormone receptor-related factors (NR1) | *MCM2, POLD1* | 0 |
| SIX1 | MA1118.1 | Homeo domain factors | HD-SINE factors | *PLK1, CKS1B* | 0 |
| Stat6 | MA0520.1 | STAT domain factors | STAT factors | *MCM2, CKS1B* | 0 |

\*Factors with p<0.05 by Mann-Whitney criterion are presented.
\*\* Profiles found in 4 or more tumor-specific promoters. Complete results of comparison are presented in Supplementary Table S1. TF recognition profiles are presented in Supplementary tables S2, S4.

**Fig. 1.**
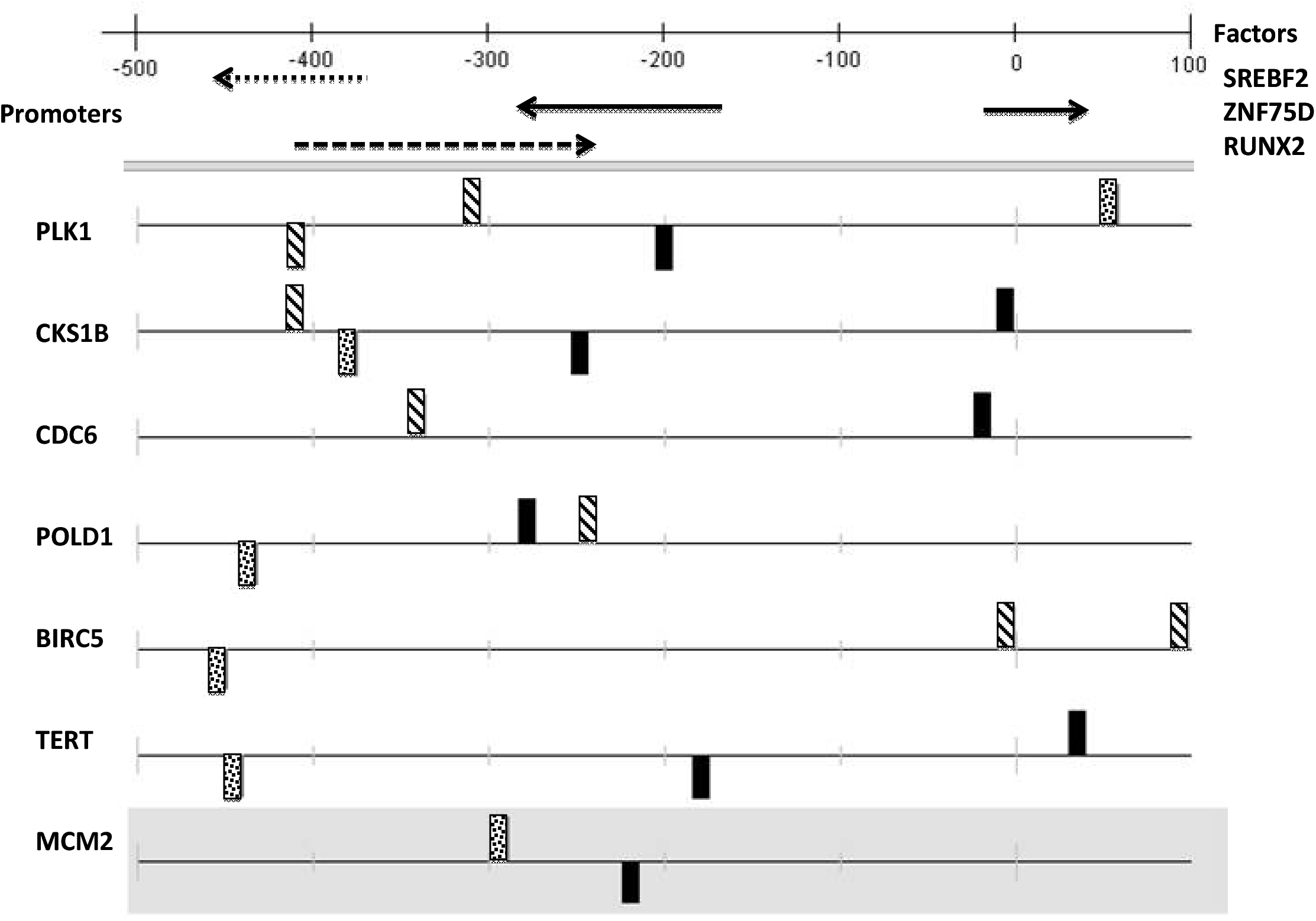
Location of the SREBF2 (dotted box), ZNF75D (black box) and RUNX2 (shaded box) recognition profiles in the promoters. Transcription start sites (0) are indicated in accordance with EPD eukaryotic promoters database.

### Interaction of transcription factors with promoters

Using the Pathway Commons resource [12], we built a network of direct regulatory interactions between the identified TFs and the products of the studied tumor-specific genes (Fig. 2). The diagram shows PCNA also interacting with the products of the genes, the promoters of which we studied. PCNA is involved in the regulation of cell proliferation and has a pronounced tumor-specific expression [15]. However, the PCNA promoter cloned by us was active in both tumor and normal cells [16]; therefore, it was initially not included in the differential analysis of promoters. In this work, the PCNA promoter (FP026732) is considered nonspecific. As seen in Fig. 2, MCM2 and PLK1 occupy a central position in this regulatory network, which is consistent with the presence of a large number of conditionally tumor-specific TF profiles in their promoters. Factors Creb3l2, CENPB, SIX1, ZNF75D, Zfx, HOXD11, and KLF11 are not participants in this regulatory network, although their recognition profiles are present in tumor-specific promoters significantly more often than in promoters of housekeeping genes. There are no data on whether these 7 TFs are involved in the regulation of the investigated promoters. However, their selection does not seem to be casual, since these TFs demonstrate differential expression or are associated with a certain prognosis in a number of tumors according to GEPIA2 (Supplementary Table S2) and, therefore, may play important roles in carcinogenesis.

**Fig. 2.**
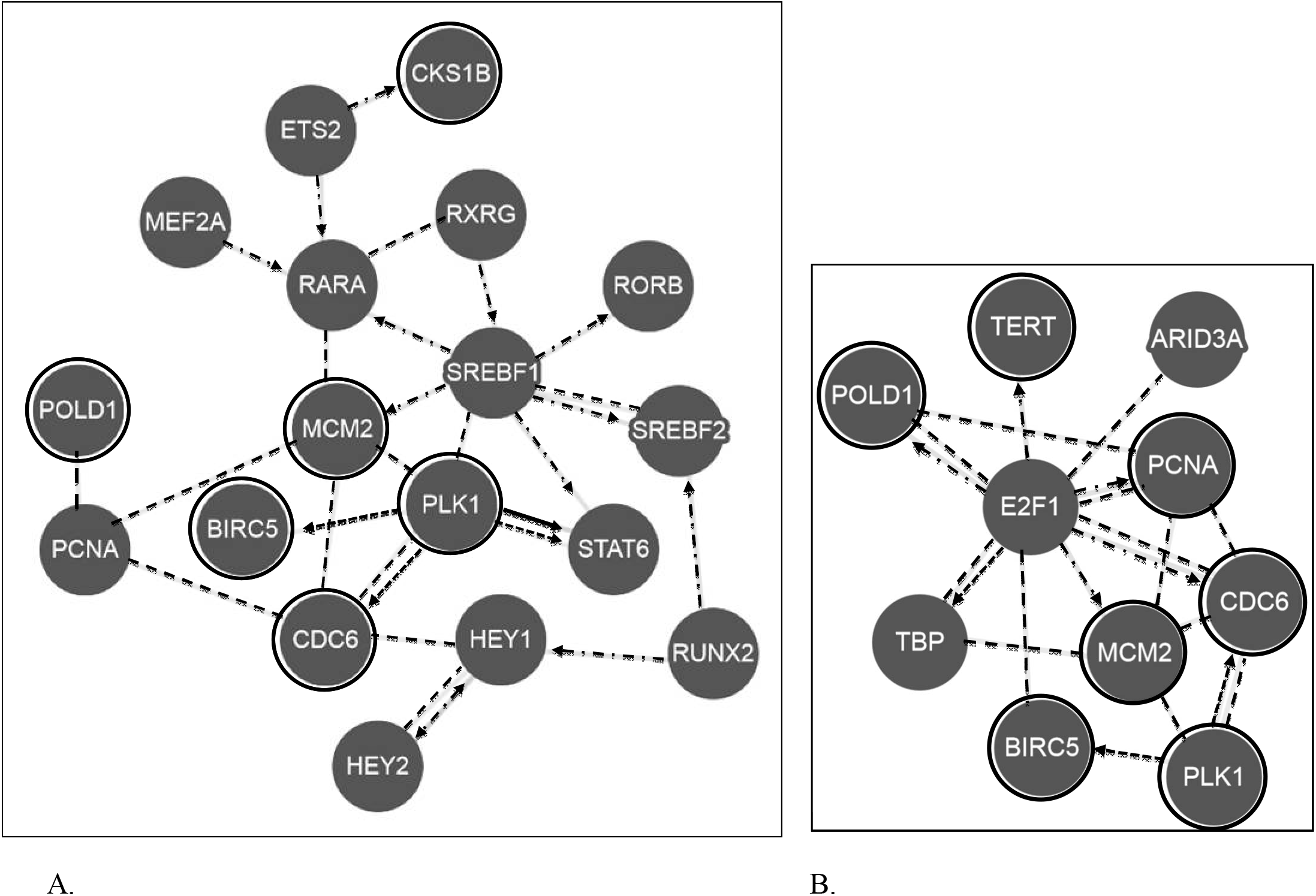
Direct regulatory interactions between the products of the studied genes (in black circles) and transcription factors by Pathway Commons. A - for conditionally tumor-specific transcription factors. B - for conditionally non-specific transcription factors. Dash-dot arrows - expression regulation; dashed arrows - modifications; solid arrow - transport control; dashed lines without arrows - binding into a complex.

### Nonspecific transcription factors

Additionally, we determined the TF profiles, which are more common in the promoters of housekeeping genes than in tumor-specific promoters (Table 3, hereinafter, conditionally nonspecific factors). It should be noted that these factors also show differential expression and may have prognostic value in some tumors (Supplementary Table S4). At the same time, five of the seven FTs of this group are involved in the development and differentiation of cells of various types (Mafb [17, 18], Arid3a [19, 20], MEIS3 [21], BHLHA15 [22, 23], BSX [24]). The other two factors are related to common mechanisms of transcription (TBP [25]) and cell cycle (E2F1 [26]). According to Pathway Commons, only TBP and E2F1 factors directly interact with the products of the studied tumor-specific genes (Fig. 2B).

**Table 3.**
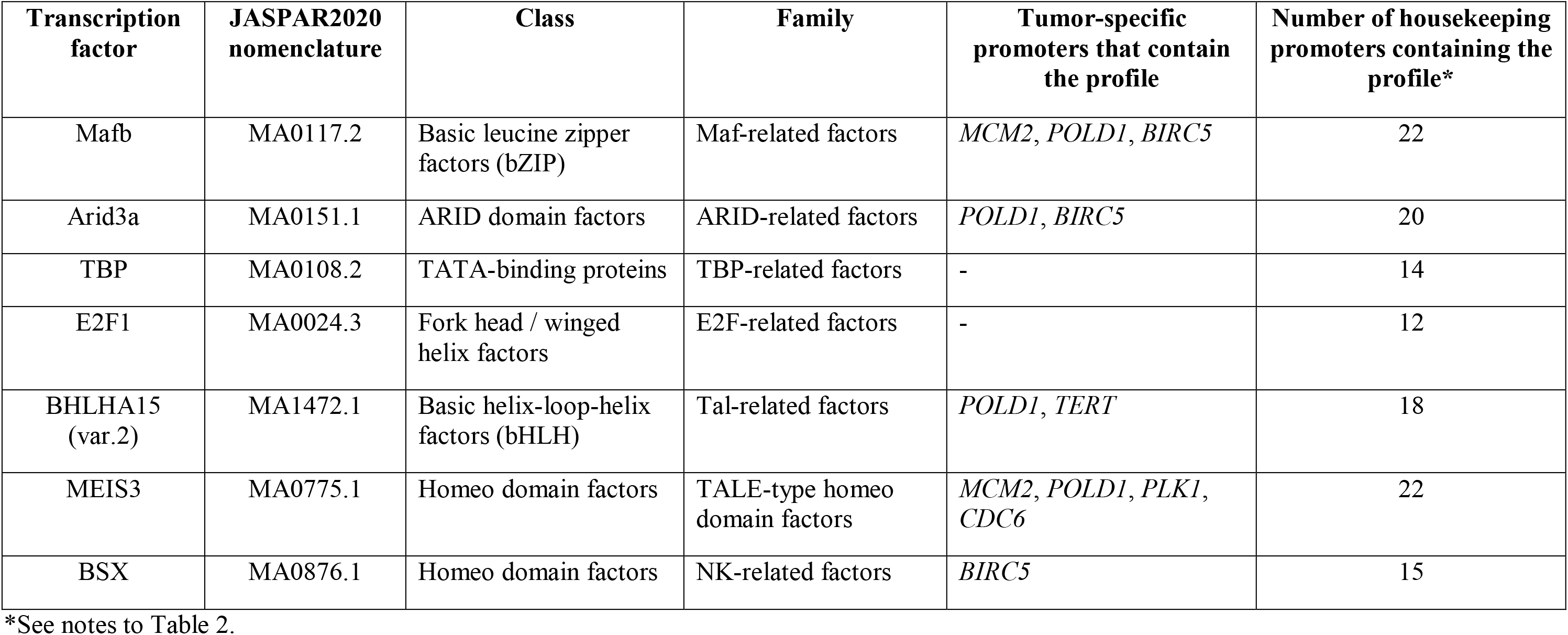
Recognition profiles of transcription factors which are more frequent in promoters of housekeeping genes than in tumor-specific promoters.*

### Analysis of chimeric promoters

To test the possible importance of the identified TFs further, we examined a set of chimeric promoters which were more active in the epidermoid carcinoma A431 cells than in normal fibroblasts (Fig. 3). These promoters were selected from the random chimeric promoter library that was constructed from fragmented tumor-specific promoters in our lab [9]. Using CiiiDER, we searched for conditionally tumor-specific and nonspecific TFs profiles in chimeric promoters, as well as in nonspecific CMV and PCNA promoters. Profiles Creb3l2, ETS2, HEY1, HEY2, and SREBF1 (var. 2) were found only in chimeric promoters which demonstrated increased activity in A431 cells (CH2, CH20, CH26, Fig. 3). These profiles were absent in nonspecific CH10 (chimeric), CMV, PCNA promoters (Supplementary Table S5). All the profiles of conditionally non-specific TFs prevailing in the promoters of the housekeeping genes (Table 3) were found in at least one of the three nonspecific promoters (Supplementary Table S6). Although this observation can hardly be estimated statistically, we tend to consider it as a trend that that supports our conclusions but requires additional verification.

**Fig. 3.**
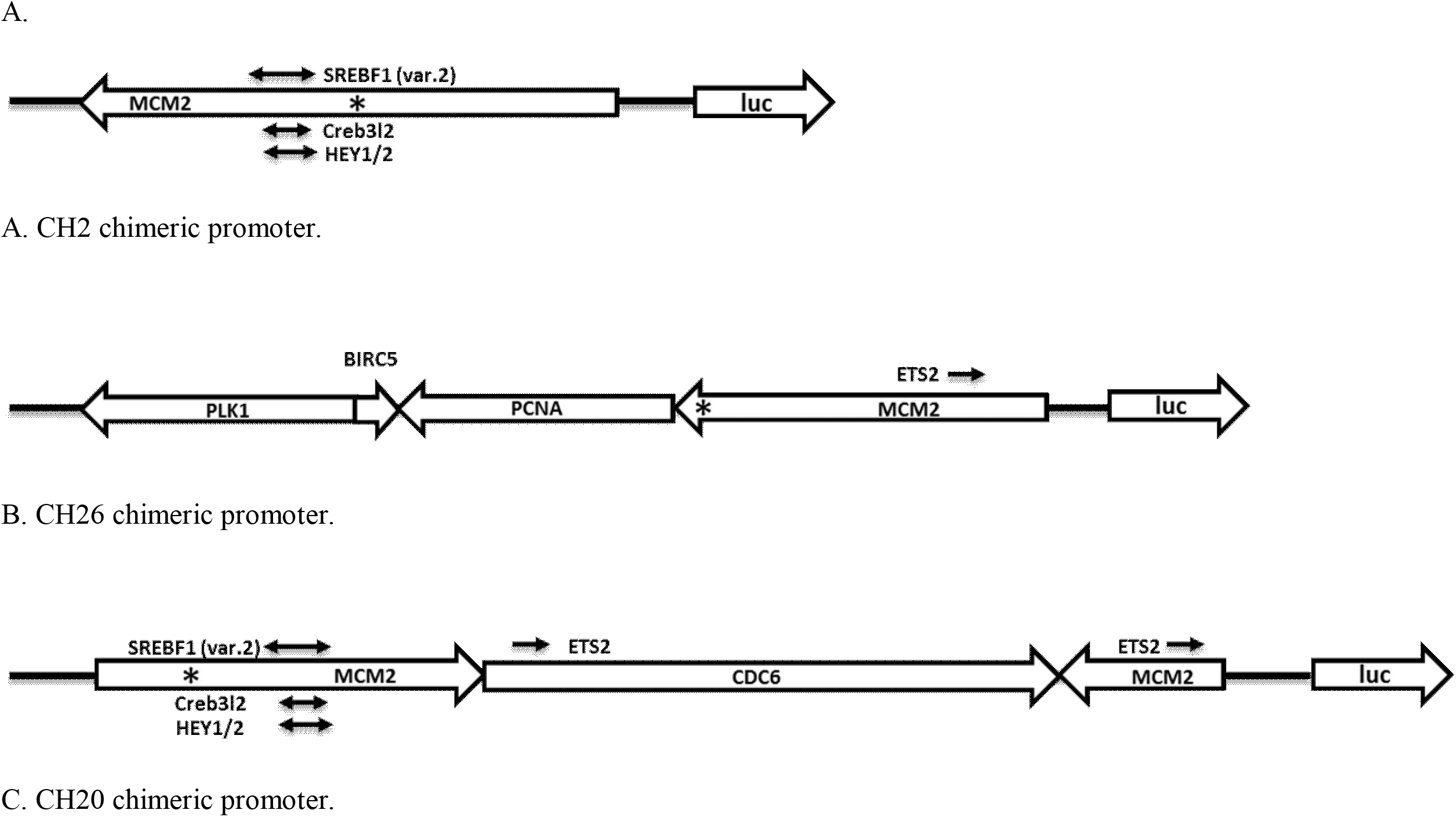
Examples of chimeric promoters (CH2, CH26, CH20) consisting of fragments of several promoters in different orientations [9]. The start sites of transcription (*) are shown as well as the location of the recognition sites for individual transcription factors.

## THE DISCUSSION

It is the primary DNA structure that determines the binding of proteins of the transcriptional apparatus and transcription factors to promoters and other regulatory elements of the genome, and through this, all the following levels of expression regulation: close and distant interactions of regulatory elements, topological units of transcription, epigenetic regulation, etc. [27].

Currently, the accumulated data on the structure of promoters already open up great prospects for the use of promoters in practical medicine, for example, as parts of genetically engineered vectors for gene therapy of tumors [28–30]. Clinical studies of gene therapy constructs containing viral and human promoters have been completed (ClinicalTrials.gov Identifier: NCT01455259; NCT00891748; NCT00197522; NCT00051480; see reviews [28, 31] and others). However, the study of the properties of promoters is far from the limit. In addition to gene therapy for cancer, natural promoters can be used in genetically engineered vectors for the treatment of other diseases, for use in industry and various biotechnological processes. Approaches with the construction of hybrid promoters [7, 13, 32–34] and chimeric promoters [9] are being developed. Therefore, the study of the primary structure of promoters, in particular, tumor-specific promoters, remains relevant.

In this work, we investigated seven tumor-specific promoters (Table 1) by the content of recognition profiles of known transcription factors. Despite the fact that both DNA sequences adjacent to TSS and distant ones are involved in the regulation of transcription, it has been shown that certain regulatory functions are retained in relatively short proximal promoters, which are minimal (core) promoters with adjacent cis- regulatory elements with a total length of several hundred nucleotides [35–37]. This is also true for the investigated promoters, in which many important regulatory elements are concentrated in the region up to –500 bp from TSS [8, 13, 14]. Therefore, we selected DNA sequences with coordinates [-499; 100] relative to TSS from the EPD database of eukaryotic promoters [10] which correspond to the previously investigated promoters [7, 8] and compared them with the promoters of housekeeping genes.

In the course of comparison, we identified 17 recognition profiles of TFs which are more frequent in tumor-specific promoters than in promoters of housekeeping genes (conditionally tumor-specific TFs, Table 2). Supplementary Table S2 additionally shows the recognition profiles of these factors in accordance with the JASPAR2020 database [5]. More than half of the tumor-specific promoters contained the profiles of the factors SREBF2, ZNF75D, Zfx (each in six promoters), RUNX2 and ETS2 (each in five promoters), and Creb3l2 (in four promoters). We also determined the preferential location zones of recognition profiles of the factors SREBF2, ZNF75D and RUNX2 in tumor-specific promoters (Fig. 1). It should be noted that the CKS1B and PLK1 promoters, which contain 10 and 9 TF profiles of this group respectively, previously showed the highest tumor specificity compared to other promoters [8].

Several tumor-specific promoters contain recognition profiles of factors, the participation of which in the regulation of these promoters is unstudied. Factors that are not involved in the first-order regulatory network (Fig. 2A) with the products of the studied tumor-specific genes include Creb3l2, CENPB, SIX1, ZNF75D, HOXD11, Zfx, KLF11. Perhaps our results will serve as an argument in favor of studying the role of these TFs in the regulation of corresponding genes.

We also identified 7 profiles of TFs, more represented in the promoters of housekeeping genes than in tumor-specific promoters (Table 3, Supplementary Table S4). Five of the seven TFs of this group are involved in the development and differentiation of cells of various types (Mafb [17, 18], Arid3a [19, 20], Meis3 [21], BHLHA15 [22, 23], BSX [24]). Normal differentiation processes are usually impaired in tumor cells [38, 39]. Cell differentiation is often considered as the reverse process of malignancy, and it has also been suggested to use it for tumor therapy [40, 41]. These observations indicate the non-casual selection of TFs on the basis of non-specificity in this case. At the same time, the differential expression of TF in this group and their association with the prognosis in some tumors (Supplementary Table S4) indicate the need for a deeper study of this issue.

It is obvious that the binding of a transcription factor to DNA depends not only on the nucleotide sequence, but also on many other parameters – the orientation of the profile, adjacent sequences, the interaction of a given TF with other regulatory molecules and sequences, etc. [35]. In addition, conditionally tumor-specific TFs may be related not so much to tumor transformation of cells as to the proliferative functions of genes, and these relations should be studied in each specific case. Nevertheless, the data obtained make it possible to investigate the regulation of promoters in direct experiments more accurately.

The study of TF recognition profiles in chimeric promoters consisting of fragments of tumor-specific promoters may serve as a test for our conclusions partly. Chimeric promoters were selected for their higher activity in A431 epidermoid carcinoma cells as compared to normal fibroblasts [9]. Five of 17 profiles of conditionally tumor-specific factors, namely the profiles Creb3l2, ETS2, HEY1, HEY2, and SREBF1 (var. 2), were selected in chimeric promoters based on tumor specificity. These profiles are not contained in the non-specific CMV, PCNA, and CH10 chimeric promoters (Supplementary Table S5). As seen in Fig. 3, the recognition profiles of the factors SREBF1 (var. 2), Creb3l2, and HEY1 / 2, located near the TSS of MCM2 promoter overlap. This may indicate competitive participation of these TFs in the regulation of transcription of chimeric promoters and the natural MCM2 gene. At the same time, all TF profiles which are more frequent in the promoters of housekeeping genes are found in at least one of the three nonspecific promoters (Supplementary Table S6). We consider this observation as a trend that supports our conclusions but requires additional verification.

This work shows that the investigated tumor-specific promoters differ from nonspecific promoters of housekeeping genes by the presence of recognition profiles for 17 transcription factors, which can be promising objects for studying regulation of cell proliferation and tumor-specific gene expression. The presence of these transcription factors in any unknown promoter may indicate tumor specificity. We believe that our results may contribute to the selection of promising natural promoters for use in genetically engineered antitumor constructs. For example, three of the seven promoters studied, namely MCM2, CKS1B, and PLK1 promoters, are more saturated with the profiles of conditionally tumor-specific TFs, which is consistent with significant tumor specificity of these promoters [8]. Chimeric promoters obtained by us earlier [9], for example, CH2, CH20, and CH26, may also be of practical importance.

It should be emphasized that the conclusions of this work were made on the basis of a theoretical analysis of the nucleotide sequence of promoters; therefore, they concern only the presence of known TF recognition profiles in promoters. The actual role of transcription factors in tumor-specific regulation of the promoters should be investigated in direct experiments.

## Abbreviations

TF: transcription factor
TSS: transcription start site
EPD: Eucaryotic promoter database
TCGA: The Cancer Genome Atlas
THPA: The Human Protein Atlas

## Financing

This work was supported by the Russian Science Foundation (project No. 19-15-00317).

## Acknowledgments

The author is grateful to Dr. Jamie Gearing (Hudson Institute of Medical Research, Australia) for valuable advices on the CiiiDER program, as well as academician E.D. Sverdlov for significant comments on the work. The author is grateful to I.P. Chernov (IBCh RAS) for moral support during the preparation of this article.

## Compliance with ethical standards

This article does not describe any research involving humans or animals as objects.

## Additional materials

Supplementary Table S1.csv – full results of comparison (enrichment) of promoters; Supplementary Table S2.pdf – recognition profiles and clinical characteristics of TFs more frequent in tumor-specific promoters. Supplementary Table S3.pdf – list of TF profiles in tumor-specific promoters. Supplementary Table S4.pdf – recognition profiles and clinical characteristics of TFs which are more frequent in the promoters of the housekeeping genes. Supplementary Table S5.pdf – profiles of conditionally tumor-specific TFs in chimeric promoters. Supplementary Table S6.pdf – profiles of conditionally nonspecific TF in chimeric promoters.

