## Supplemental Table 2 for "Looking for Tumor Specific Transcription Factors. Study of Promoters in Silico"

Table S2. Recognition profiles of transcription factors which are more frequent in tumor-specific promoters than in promoters of housekeeping genes.*

| **Transcription factor** | **Nomenclature by JASPAR2020** | **Recognition profile by JASPAR2000** | **Tumors with differential expression (GEPIA2, TCGA)*** | **Prognostic significance (THPA)**** |
|---|---|---|---|---|
| RUNX2 | MA0511.2 | 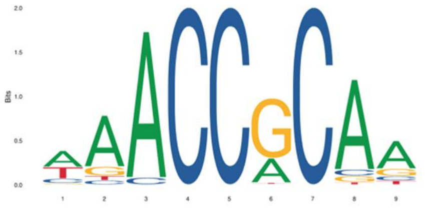 | ESCA, LAML, PAAD, STAD, THYM (+); TGCT(-) | Renal cancer (-); urothelial cancer (-), stomach cancer (-) |
| Creb3l2 | MA0608.1 | 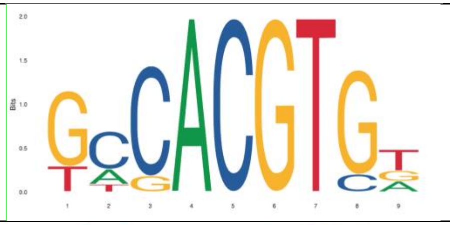 | DLBC, GBM, LGG, PAAD, SKCM, THYM (+); CESC, OV, UCEC, UCS (-) | N.P. |
| SREBF2 | MA0596.1 | 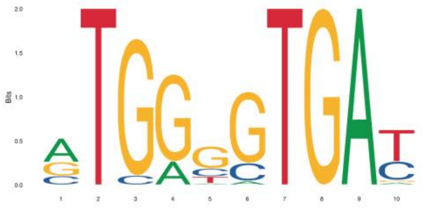 | DLBC, THYM (+); GBM, KIRC (-) | Renal cancer (+) |
| ETS2 | MA1484.1 | 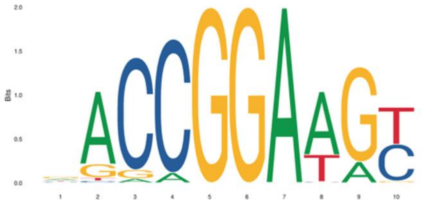 | COAD, PAAD, READ, STAD (+); BLCA, BRCA, DLBC, GBM, KICH, LUAD, LUSC, OV, PRAD, SKCM, THCA, UCEC, USC (-) | N.P. |
| CENPB | MA0637.1 | 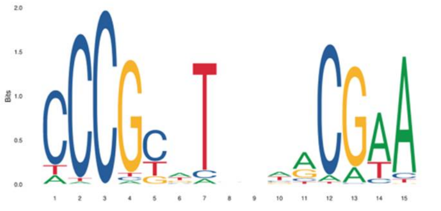 | DBLC, GBM, LGG, PAAD, THYM (+) | Liver cancer (-) |

| | | | | |
|---|---|---|---|---|
| HEY1 | MA0823.1 | 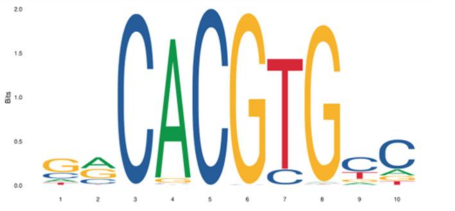 | GBM, KICH, KIRC, LGG, PAAD, SKCM, THYM, UCS (+) BRCA, LAML, LUAD (-) | N.P. |
| HEY2 | MA0649.1 | 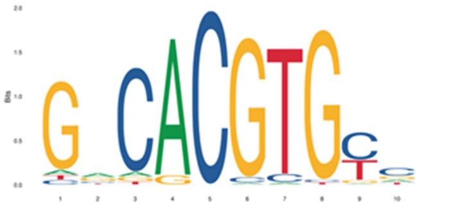 | GBM, KIRC, LGG, THCA, THYM (+); BRCA, CESC, TGCT (-) | N.P. |
| RARA::RXRG | MA1149.1 | 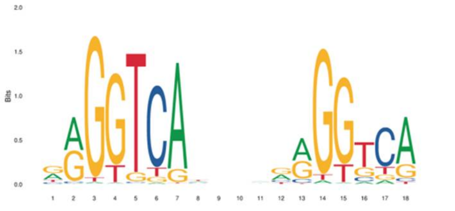 | RARA— ESCA, GBM, LAML, LGG, PAAD, STAD (+); CESC, DLBC, LUSC, TGCT, THYM (-); RARG — SKCM, THCA (+); BRCA, CESC, COAD, GBM, LUAD, LUSC, READ (-) | RARA – endometrial cancer (+); RXRG – thyroid cancer (+) |
| Srebf1(var.2) | MA0829.1 | 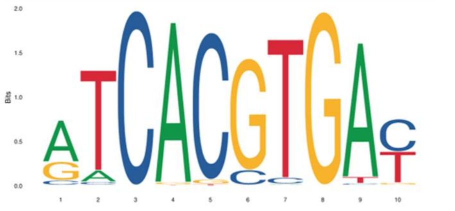 | DLBC, KICH, PAAD, THYM (+); ACC, OV, TGCT, UCS (-) | endometrial cancer (+), pancreatic cancer (+) |
| ZNF75D | MA1601.1 | 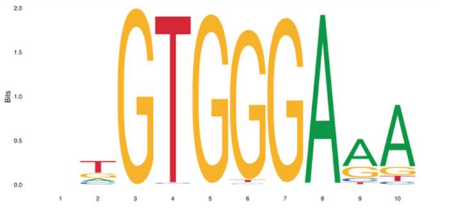 | LAML, THYM (+); LUSC, OV, USC (-) | Urothelial cancer (+), renal cancer (+) |

| | | | | |
|---|---|---|---|---|
| Zfx | MA0146.2 | 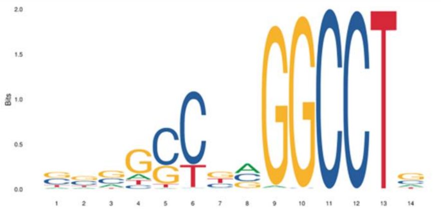 | GBM, LAML, LGG, THYM (+); UCEC, UCS (-) | N.P. |
| HOXD11 | MA0908.1 | 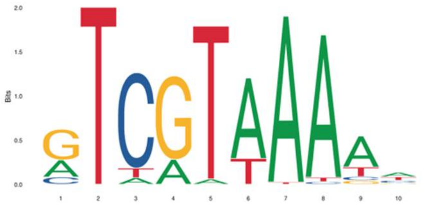 | ESCA, GBM, HNSC, LUSC (+); COAD, KICH, KIRP, PRAD, UCEC, UCS (-) | N.P. |
| KLF11 | MA1512.1 | 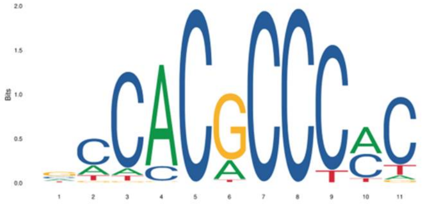 | BLCA, BRCA, SKCM, TGCT, UCEC, UCS (-) | N.P. |
| MEF2A | MA0052.4 | 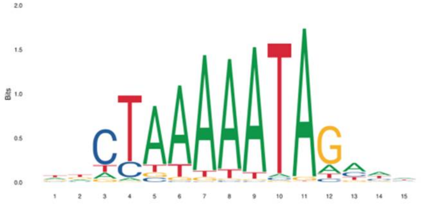 | PAAD (+); BLCA, CESC, COAD, LUAD, LUSC, READ, SKCM, UCEC, UCS (-) | Renal cancer (+), stomach cancer (-) |
| RORB | MA1150.1 | 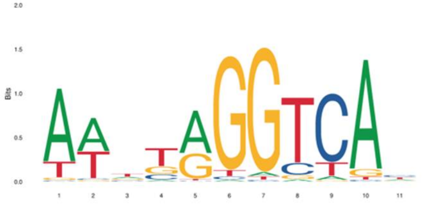 | GBM, LGG (-) | N.P. |

| | | | | |
|---|---|---|---|---|
| SIX1 | MA1118.1 | 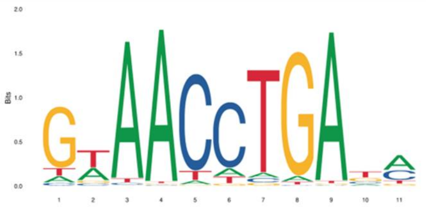 | BLCA, BRCA, CESC, ESCA, GBM, KICH, LGG, LUAD, LUSC, OV, THYM, UCEC, UCS (+); TGCT, THCA (-) | Endometrial cancer (-), lung cancer (+) |
| Stat6 | MA0520.1 | 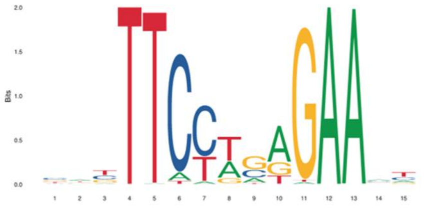 | LAML, PAAD (+); CESC, LGG, LUAD, LUSC, OV, PRAD, UCEC, UCS (-) | Urothelial cancer (+), endometrial cancer (+), thyroid cancer (+) |

*Tumor nomenclature by The Cancer Genome Atlas. (+) – enhanced expression; (-) – decreased expression.

** Prognostic significance according to The Human Protein Atlas. (+) favorable marker; (-) unfavoreable marker. N.P. – not prognostic.
