## Supplemental Table 3 for "Looking for Tumor Specific Transcription Factors. Study of Promoters in Silico"

Table S3. Recognition profiles of transcription factors that are characteristic to tumor-specific promoters.

| Promoter | Transcription factor recognition profiles |
|---|---|
| *MCM2* | Creb3l2, SREBF2, ETS2, CENPB, HEY1, HEY2, RARA::RXRG, SREBF1(var2), ZNF75D, Zfx, MEF2A, RORB, Stat6 |
| *CKS1B* | RUNX2, SREBF2, ETS2, CENPB, RARA::RXRG, ZNF75D, Zfx, HOXD1, SIX1, Stat6 |
| *PLK1* | RUNX2, Creb3l2, SREBF2, ETS2, HEY1, HEY2, SREBF1(var.2), ZNF75D, SIX1 |
| *POLD1* | RUNX2, SREBF2, RARA::RXRG, ZNF75D, Zfx, HOXD11, KLF11, RORB |
| *TERT* | Creb3l2, SREBF2, ETS2, HEY1, HEY2, ZNF75D, Zfx |
| *CDC6* | RUNX2, Creb3l2, ETS2, CENPB, ZNF75D, Zfx |
| *BIRC5* | RUNX2, SREBF2, SREBF1(var.2), Zfx, KLF11, MEF2A |
