## Supplemental Table 4 for "Looking for Tumor Specific Transcription Factors. Study of Promoters in Silico"

Table S4. Recognition profiles of transcription factors which are more frequent in promoters of housekeeping genes than in tumor-specific promoters *

| **Transcription factor** | **Nomenclature by JASPAR2020** | **Recognition profile by JASPAR2000** | **Tumors with differential expression (GEPIA2, TCGA)*** | **Prognostic significance (THPA)**** |
|---|---|---|---|---|
| Mafb | MA0117.2 | 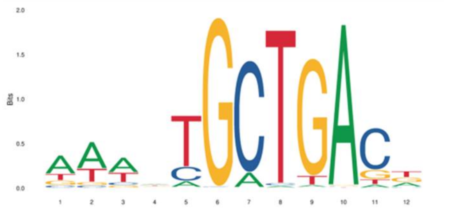 | GBM, LGG, OV, PAAD, STAD, TGCT, UCS (+); SKCM (-) | Endometrial cancer (-) |
| Arid3a | MA0151.1 | 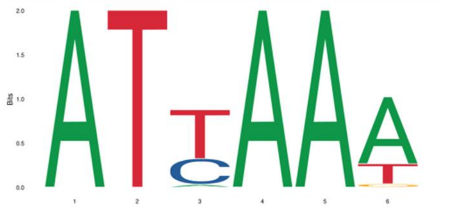 | COAD, READ, STAD (+); TGCT, THYM (-) | Pancreatic cancer (+), renal cancer (-) |
| TBP | MA0108.2 | 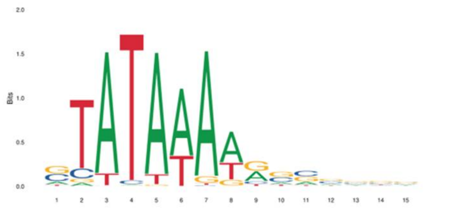 | DLBC, THYM (+); LAML, TGCT (-) | Liver cancer (-) |
| E2F1 | MA0024.3 | 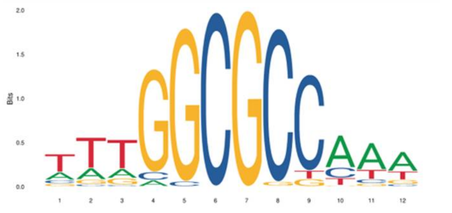 | ACC, BLCA, BRCA, CESC, COAD, DLBC, ESCA, GBM, HNSC, KICH, KIRC, KIRP, LIHC, LUAD, LUSC, OV, PAAD, READ, SKCM, STAD, THCA, THYM, UCEC, UCS (+) | Cervical (+), liver (-), endometrial (-), renal (-), thyroid (+), pancreatic (-) cancer |

| | | | | |
|---|---|---|---|---|
| BHLHA15(var.2) | MA1472.1 | 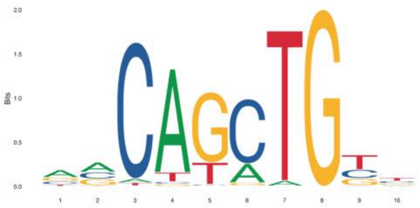 | OV, PRAD (+); PAAD, STAD (-) | N.P. |
| MEIS3 | MA0775.1 | 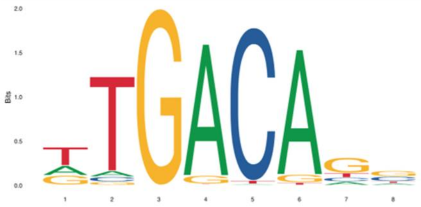 | DLBC, HNSC, KIRP, PAAD (+); CESC, COAD, OV, READ, THCA, UCEC, UCS (-) | Renal cancer (-) |
| BSX | MA0876.1 | 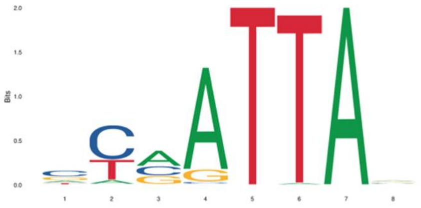 | — | N.P. |

*Tumor nomenclature by The Cancer Genome Atlas. (+) – enhanced expression; (-) – decreased expression.

** Prognostic significance according to The Human Protein Atlas. (+) Favorable marker; (-) unfavoreable marker. N.P. – not prognostic.
