## Supplemental Table 5 for "Looking for Tumor Specific Transcription Factors. Study of Promoters in Silico"

Table S5. Recognition profiles of of conditionally tumor-specific TFs in chimeric promoters.

| **Transcription factor** | **JASPAR2020 nomenclature** | **Promoters** | | | | | | | | | |
|---|---|---|---|---|---|---|---|---|---|---|---|
| | | **CH2** | **CH8** | **CH9** | **CH12** | **CH16** | **CH20** | **CH26** | **CH10** | **CMV** | ***PCNA*** |
| RUNX2 | MA0511.2 | - | - | - | - | + | - | - | - | + | - |
| Creb3l2* | MA0608.1 | + | - | - | - | - | + | - | - | - | - |
| SREBF2 | MA0596.1 | - | + | - | - | + | - | - | + | - | - |
| ETS2* | MA1484.1 | + | - | - | - | - | + | + | - | - | - |
| CENPB | MA0637.1 | + | + | + | + | - | + | + | + | - | - |
| HEY1* | MA0823.1 | + | - | - | - | - | + | - | - | - | - |
| HEY2* | MA0649.1 | + | - | - | - | - | + | - | - | - | - |
| RARA::RXRG | MA1149.1 | - | + | + | + | + | - | - | + | + | - |
| Srebf1(var.2)* | MA0829.1 | + | - | - | - | - | + | - | - | - | - |
| ZNF75D | MA1601.1 | - | - | - | + | + | + | - | + | + | + |
| Zfx | MA0146.2 | - | - | - | - | + | + | - | + | - | + |
| HOXD11 | MA0908.1 | - | - | - | - | - | - | - | - | - | + |
| KLF11 | MA1512.1 | - | - | - | - | + | - | - | - | - | - |
| MEF2A | MA0052.4 | - | - | - | - | - | - | - | - | - | - |
| RORB | MA1150.1 | - | - | - | - | + | - | - | + | - | - |
| SIX1 | MA1118.1 | - | - | - | - | - | - | - | - | - | - |
| Stat6 | MA0520.1 | - | - | - | - | - | - | - | - | - | - |

* Location of these profiles in promoters is shown of Fig. 3.
