## Supplemental Table 6 for "Looking for Tumor Specific Transcription Factors. Study of Promoters in Silico"

Table S6. Recognition profiles of conditionally non-specific transcription factors in chimeric promoters

| **Transcription factor** | **JASPAR2020 nomenclature** | **Promoters** | | | | | | | | | |
|---|---|---|---|---|---|---|---|---|---|---|---|
| | | **CH2** | **CH8** | **CH9** | **CH12** | **CH16** | **CH20** | **CH26** | **CH10** | **CMV** | ***PCNA*** |
| Mafb | MA0117.2 | - | - | - | - | + | - | - | + | + | + |
| Arid3a | MA0151.1 | - | + | - | - | + | - | - | - | + | + |
| TBP | MA0108.2 | - | - | - | - | - | - | - | - | + | + |
| E2F1 | MA0024.3 | - | - | - | - | - | - | - | - | - | - |
| BHLHA15 (var.2) | MA1472.1 | - | - | - | - | - | - | - | - | - | + |
| MEIS3 | MA0775.1 | - | - | - | - | + | - | - | - | - | + |
| BSX | MA0876.1 | - | - | - | - | + | - | - | - | + | + |
